# Structural Insights into the Interaction Between Adenovirus C5 Hexon and Human Lactoferrin

**DOI:** 10.1101/2023.10.09.561496

**Authors:** Arun Dhillon, B. David Persson, Alexander N. Volkov, Hagen Sülzen, Alan Kádek, Petr Pompach, Sami Kereïche, Martin Lepšík, Katarina Danskog, Charlotte Uetrecht, Niklas Arnberg, Sebastian Zoll

**Author notes:** Corresponding author: SZ.

## Abstract

Adenovirus (AdV) infection of the respiratory epithelium is common, but poorly understood. Human AdV species C types, such as HAdV-C5, utilize the Coxsackie-adenovirus receptor (CAR) for attachment and subsequently integrins for entry. CAR and integrins are however located deep within the tight junctions in the mucosa where they would not be easily accessible. Recently, a model for CAR-independent AdV entry was proposed. In this model, human lactoferrin (hLF), an innate immune protein, aids the viral uptake into epithelial cells by mediating interactions between the major capsid protein, hexon, and yet unknown host cellular receptor(s). However, a detailed understanding of the molecular interactions driving this mechanism is lacking. Here, we present a new cryo-EM structure of HAdV-5C hexon at high resolution alongside a hybrid structure of HAdV-5C hexon complexed with human lactoferrin (hLF). These structures reveal the molecular determinants of the interaction between hLF and HAdV-C5 hexon. hLF engages hexon primarily via its N-terminal lactoferricin (Lfcin) region, interacting with hexon’s hypervariable region 1 (HVR-1). Mutational analyses pinpoint critical Lfcin contacts and also identify additional regions within hLF that critically contribute to hexon binding. Our study sheds more light on the intricate mechanism by which HAdV-C5 utilizes soluble hLF/Lfcin for cellular entry. These findings hold promise for advancing gene therapy applications and inform vaccine development.

## INTRODUCTION

Adenoviruses are non-enveloped viruses with a capsid of icosahedral symmetry that encapsulates a double-stranded DNA genome (1). Currently, more than 110 human adenoviruses (HAdV) types have been identified that are categorized into “species” A through G based on tissue tropism and genetic diversity (2). HAdVs usually cause self-limiting ocular (species B, C, D, E), respiratory (species B, C, E), gastrointestinal (species A, F) and urinary tract (species B) infections, which might turn lethal in immuno-compromised individuals and children (3). Species C adenoviruses that include HAdV-5 may also cause persistent infections of the adenoids and tonsils (4, 5) resulting in viral shedding for months (6). Historically, adenoviruses have played pivotal roles as molecular biology tools in the discovery of eukaryotic biological mechanisms such as RNA splicing and protein folding (7, 8). In addition, the non-integrating nature of adenoviruses has been harnessed in the field of gene therapy (9) where a major obstacle to overcome for the use of HAdVs in gene therapy is the presence of pre-existing immunity (10) and the relatively poor transduction efficiency in the target tissue (11). Nevertheless, after decades of research an HAdV based gene therapy was recently approved by regulators in the United States of America (9). In addition, vaccines based on modified adenoviruses against Ebola (12), HIV (13), ZIKV (14), RSV (15) and SARS-CoV2 (16) are currently in various phases of clinical trials or already being used in the clinic.

Species C adenoviruses infect respiratory epithelial cells by interacting with the cellular receptor CAR using the protruding fiber protein, followed by secondary interactions between the penton base and α_v_β_3_ integrins that triggers entry (1, 17, 18). This fundamentally simple model is likely used in immortalized cell lines but is highly unlikely in the polarized epithelium where CAR is hidden deep withing the tight junctions (19-21). However, additional models and mechanisms have been suggested to overcome this dilemma. HAdV-C5 infection may be facilitated by airway macrophages that take up invading HAdV particles resulting in secretion of IL-8 as part of if their inflammatory response (22). The secretion of IL-8 results in the apical migration of the CAR isoform CAR^Ex7^ and α_v_β3 from the basolateral membrane while enhancing the expression of the CAR^Ex8^ at the apical membrane (22). In addition, several CAR independent cell entry pathways have been identified for HAdV-C5, such as the use of heparin on hepatocytes (23), α_v_β_5_ on melanoma and breast cancer cells (24), and Toll-like receptor 4 on dendritic cells (25). Finally, several different types of HAdVs have been shown to use different soluble components present in body fluids enhance the infection. For example, HAdV-5C has been shown to recruit and hijack coagulation factor X (FX) for infection of hepatocytes (23), which not only enhances infection but also controls the tropism by directing the virus to the liver. During infection, FX binds to protruding loops of the HAdV-hexon on the capsid thus allowing FX to bridge an interaction between the cellular receptor and the virus particle. Hexon is a trimeric, barrel-shaped protein that represents the main building block of the virus capsid. In addition to coagulation factors, species C HAdVs have been shown to use hLF and its proteolytic cleavage product lactoferricin (Lfcin), a highly positively charged, 49 amino acid peptide with antimicrobial activity (26), for enhanced infection of epithelial cells (27, 28). Even though this mechanism is less well described, both hLF and Lfcin have been suggested to bind the HVR-1 of hexon acting as a bridge between the virus capsid and cell surface receptor(s). Interestingly, similar mechanisms have been described for species A HAdVs that bind factor IX for a more effective infection (29, 30) indicating that this mechanism might be an efficient way for HAdVs to bypass initial infection using CAR.

In this study we present a new cryo-EM structure of HAd-5C hexon at FSC_0.143_ of 2.9 Å together with a detailed hybrid structure of hLF in complex with HAdV-5C hexon providing structural insight into the previously reported interaction between hLF and HAdV-C5 hexon (28). We show that hLF engages the HVR-1 of one hexon protomer via its N-terminal Lfcin region with additional stabilizing contacts between the hLF C-terminal lobe and a second hexon protomer. In addition, structure-based mutational studies identified key residues within Lfcin as well as an adjacent helix in the N-lobe. Taken together, our structural and biophysical characterizations shed more light onto the mechanism by which HAdV-C5 utilizes soluble hLF/Lfcin for cellular entry. These detailed insights may eventually aid the development of novel adjuvants for the use in HAdV-C5 based gene therapy.

## RESULTS

### The interaction between HAdV-C5 hexon and hLF is charge driven

Based on a series of viral transduction experiments it was previously suggested that the interaction between HAdV-C5 hexon and hLF involves the stretch of acidic amino acid residues in the HAdV-C5 hexon HVR-1 and the basic N-terminal region of hLF (28) (Fig. 1A and B). When proteolytically liberated, the N-terminal region of hLF, Lfcin, is equally capable of promoting transduction (26). Here, we provide direct biophysical evidence for a charge-driven interaction. Using size exclusion chromatography (SEC) and native mass spectroscopy (MS) we could show that stability of the complex is highly dependent on the ionic strength of the buffer used which is typical for charge-driven interactions (Fig. 1C and D). In SEC the HAdV-C5 hexon:hLF complex does stay intact only at a salt concentration of ≤ 5 mM (Fig. 1C). Any further increase in salt concentration leads to gradual disruption of the complex. At 10 mM NaCl the complex largely disintegrates with free hLF adsorbing to the matrix of the SEC column. Therefore, at 10 mM NaCl, no additional peak corresponding to free hLF such as in the presence of 150 mM NaCl, can be observed. Excess hLF not bound to hexon could afterwards be eluted from the column with a higher salt concentration. To rule out aberrant behavior of the protein due to nonspecific interaction with the stationary phase of the column under low salt conditions we carried out native MS measurements. Consistent with observations made in SEC, binding of hLF to hexon could only be observed under low ionic strength conditions (Fig. 1D, Fig. S1). In this case ammonium acetate (AA) was used as an MS-compatible buffer surrogate. In the presence of 20 mM AA the hexon:hLF complex was not stable. Compared to free hexon, only a fraction of hexon:hLF at 3:1 (hexon protomers:hLF) stoichiometry was detectable. Upon dilution from 20 mM AA to around 5 mM predominantly a complex with 3:1 stoichiometry was observed with smaller amounts of a complex with 2 hLF molecules per hexon trimer as well as free hexon.

**Figure 1.**
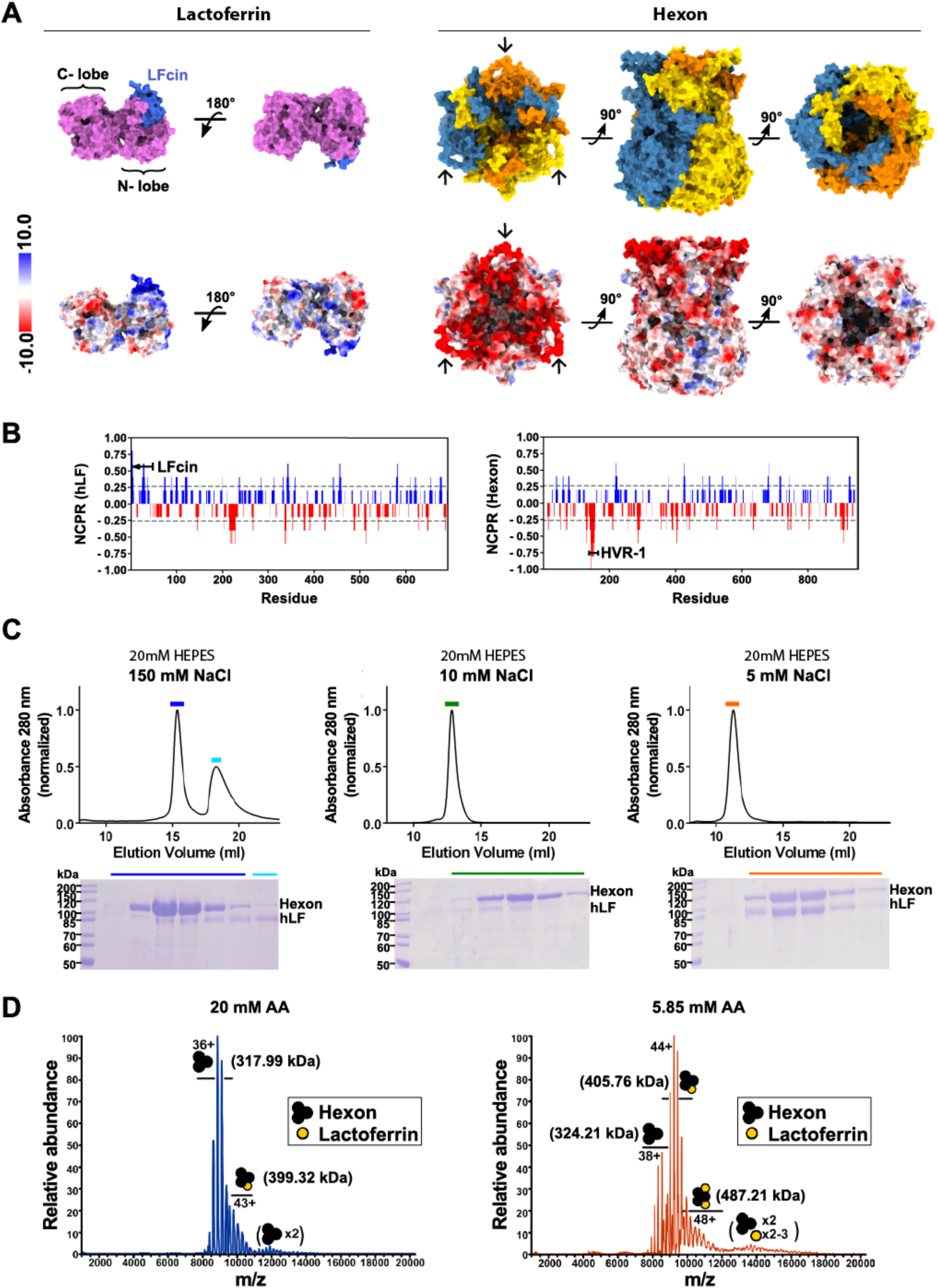
Effect of ionic strength on hexon:hLF complex formation. (A) Surface views of Lactoferrin (PDB: 1LFG) (left) and hexon, completed with AlphaFold2 (see methods) (right). The Lfcin peptide on the N-terminal lobe of LF is colored in blue. The three protomers of hexon are shown in different colors: orange, yellow, blue. The HVR-1 loop of each subunit is marked with an arrow. Surface representations coloured according to electrostatic potential are depicted below. Notably, the area on Lactoferrin corresponding to the Lfcin peptide exhibits a strong positive charge while the HVR-1 loops of hexon are strongly negatively charged. (B) Net charge per residue (NCPR) distribution along the amino acid sequence of hLF (left) and hexon (right). Blue color indicates positive NCPR and red colour indicates negative NCPR. Residues corresponding to the Lfcin and HVR-1 have been highlighted. NCPR distribution was calculated via CIDER with the default blob size. (C) Hexon:hLF complex formation at different ionic strengths during size exclusion chromatography (top). Reducing SDS gels (bottom) confirm the co-elution of hexon and hLF. Fractions loaded onto the SDS gel have been highlighted with colored bars in the corresponding chromatogram (D) Native MS of 0.5 µM hexon:hLF complex electro-sprayed from 20 mM (left) and 5.85 mM (right) ammonium acetate (AA) pH 7.0. Hexon is detected exclusively trimeric (black) with up to 2 hLF molecules bound (yellow). Main charge states and corresponding molecular weights are indicated. Small amounts dimeric hexon trimer are artefacts due to ESI concentration effects. For more detailed peak assignment, see Fig. S1.

### Cryo-EM structure of HAdV-C5 hexon at 2.9 Å

The structure of HAdV-C5 hexon was reconstructed to FSC_0.143_ of 2.9 Å (Fig. 2A, B and Table S1). To our knowledge this is the highest resolution structure of HAdV-C5 hexon determined by cryo-EM. Due to high stability of the trimer the resolution is relatively uniform across the whole structure and thus it could be modelled unambiguously with a high average cross-correlation of 0.736 (Fig. 2C). Throughout the length of hexon, the protomers are in close contact forming large interfaces in addition to intertwined regions at the ‘turrets’ and the base. Each protomer contributes approximately 16 000 Å^2^ to the interface with the two other protomers. Significantly lower resolution is observed only for the hypervariable loop regions, the ‘turrets’ at the top of the molecule which are largely disordered and could not be built due to a lack of traceable density in the final reconstruction. Specifically, regions 137-164, 188-190, 252-256, 271-278, 431-436 could not be sufficiently reconstructed. These regions are also lacking in the crystal structure (3TG7) of hexon. The same structure was used a starting model to build hexon in the cryo-EM reconstruction, but no major conformational changes were observed. The RMSD between both structures is 0.9 Å, indicating high overall similarity (Fig. S2).

**Figure 2.**
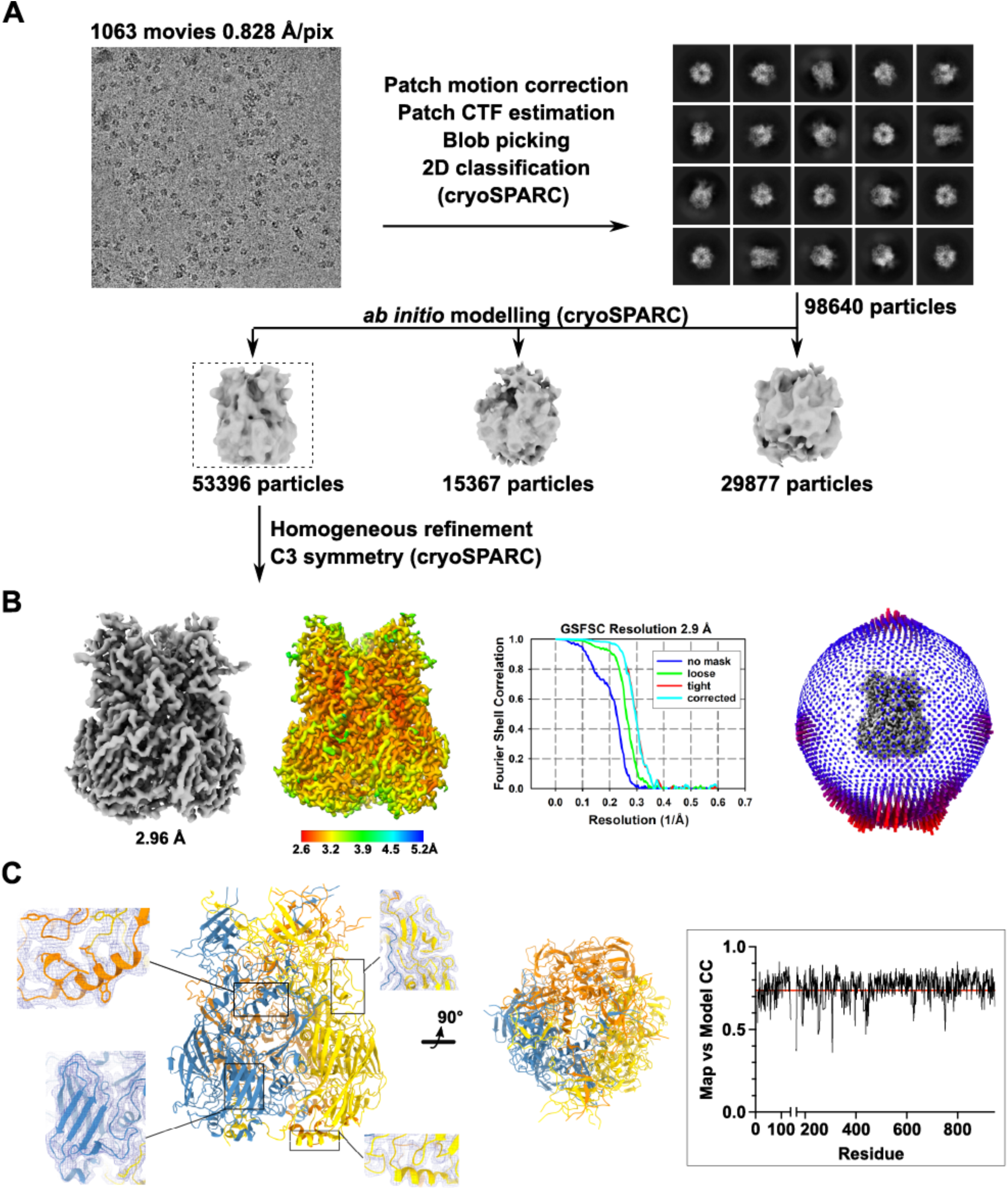
Structure of HAdV-C5 hexon determined by single particle cryo-EM at 2.9Å. (A) Visual representation of the work flow adopted for data processing using cryoSPARC. A particle set identified by iterative, reference free 2D classification was used to obtain 3 *ab initio* models. The model with the most recognizable features was selected (boxed) and homogeneously refined to reconstruct the trimeric hexon. 53396 particles contributed to the final reconstruction. A representative micrograph (top left) is shown alongside a panel of selected 2D class averages. (B) The final, unsharpened map of the hexon reconstruction is shown in grey and colored according to local resolution. Next to the gold-standard FSC curves is angular distribution plot of the reconstruction (C) Side and bottom view of the hexon model built from the cryo-EM reconstruction. The three protomers of the hexon are depicted in the color scheme used previously. Insets show close-up views of the model inside the unsharpened map from different regions of the hexon. The plot of the Map vs Model cross-correlation for one protomer (boxed) shows a high average cross-correlation of 0.736 (red line) at FSC_0.143_: 2.9 Å.

### Integrative modelling of the HAdV-C5 hexon:hLF complex

Despite many attempts, strong preferential orientation of the hexon:hLF complex on EM grids could not be overcome. Unlike for hexon alone, which also showed a higher number of top and bottom views (Fig. 2B), this phenomenon was exacerbated in the complex and precluded its reconstruction. Standard remedial approaches such as stage tilting and the use of detergents were ineffective. Furthermore, grid preparation was highly irreproducible with only small percentages of intact complex particles being detectable. Good 2D class averages were however obtained for top and bottom views of the complex. We therefore set out to use this information in calculation of a hybrid atomic model of the complex, adapting a protocol by Walti et al (31). Implemented in the program XPLOR-NIH (32), this protocol allows fitting of protein models to 2-dimensional electron density projection images. In the first step of our integrative modeling approach, we performed crosslinking mass spectrometry (XL-MS) with the HAdV-C5 hexon:hLF complex using the homobifunctional crosslinker disuccinimidyl dibutyric urea (DSBU). 2 pairs of crosslinks were identified (Fig. 3A and Fig S3). The hexon residues (S174, S446) crosslinked to hLF residues are present in the ‘turrets’, while one of the corresponding hLF residues is located in the N-lobe (K282) and the second (K676) one at the junction of N- and C-lobe. Since, hexon is a homotrimer, in principle, hLF could be crosslinked to residues belonging to the same protomer or to residues on different protomers of hexon. However, only when hLF was assumed to be crosslinked to the same hexon protomer did the Lfcin region face hexon. As prior evidence exists for involvement of Lfcin in hexon binding (28) it was assumed that hLF is crosslinked to residues on same protomer of hexon (hexon S446: hLF K282; hexon S174: hLF K676). Next, we carried out a molecular dynamics simulation of hexon alone (Fig. 3A) and selected a conformation from the latter part of the trajectory, post-convergence of RMSD, that closely mirrors the observed hexon conformation in experimental 2D classes of the complex (Fig. 3B and Fig. S4). Complex projections suggest a conformational change in the flexible ‘turret’ region of hexon upon binding of hLF.

**Figure 3.**
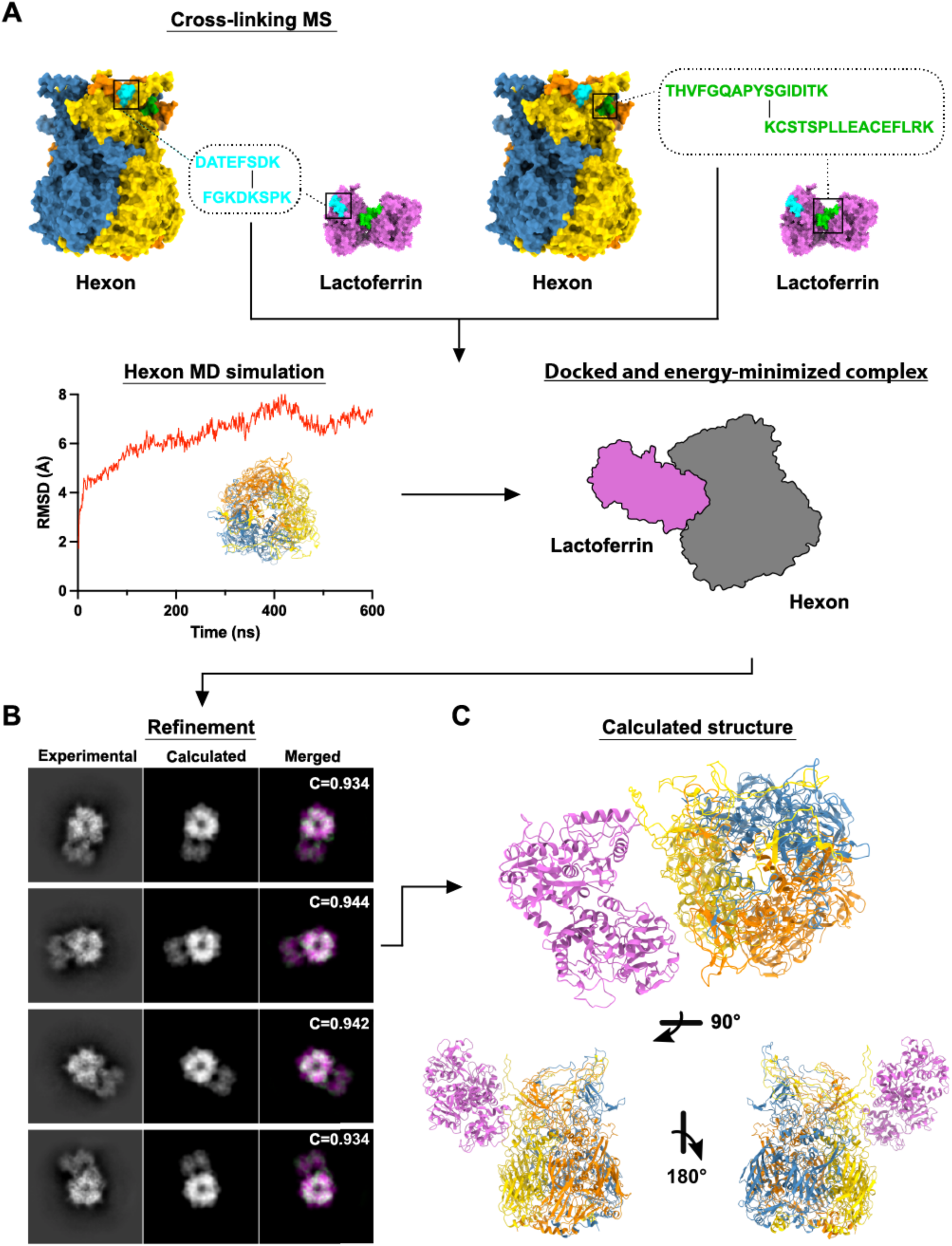
Visual representation of computation workflow for a hybrid model of the hexon:hLF complex. (A) Top, Cross-linking MS revealed two cross-links, locations of identified peptides are highlighted on surface representations of hLF (PDB: 1LFG) and hexon in cyan and green. Peptide sequences are shown in the corresponding color, crosslinked residues are marked with a black line. Bottom left; Hexon MD simulation – the evolution of backbone RMSD, overlay of several snapshots from the MD trajectory. Bottom right; Identified cross-links were used as distance restrains in the subsequent docking and energy minimization of the complex in XPLOR-NIH, visualized by representative schematic (B) Refinement of the docked hexon:hLF complex against experimental 2D class averages of the complex using XPLOR-NIH (see methods). The left column shows the experimentally obtained 2D class averages for the hexon:hLF, the central column shows the calculated back projections for the hexon:hLF complex obtained after docking and energy minimization, and the right column shows the result of superposition of experimental (green) and calculated (magenta) projections. Cross-correlation (C) of experimental and calculated projections is shown for individual projections (D) Different views of the final hybrid model (best C-value) of hexon:hLF complex. LF is colored pink, protomers of hexon are individually colored using the color scheme as before.

These steps created the hexon input model and established the principal binding site and distance constrains for docking of the complex in XPLOR-NIH. The docked and energy-minimized complex (Fig. 3A) was then rigid body refined against the 2D projections and models were calculated (Fig 3B and C). Models without crosslink distance violations were ranked according to their correlation with the experimental projections of the complex (Fig. 3B and Fig S5). The model of HAdV-C5 hexon:hLF complex with the highest C-value was selected as the final model and used for further analysis (Fig. 3C). In the selected model residues 1-49 of hLF, constituting Lfcin, are positioned facing the hexon HVR-1, while another previously not reported point of close contact is near crosslink 2 (hexon S174-hLF K676) between the C-lobe of hLF and an adjacent protomer within hexon.

### Residues on both hLF lobes contribute to HAdV-C5 hexon binding

In order to validate our model and obtain further molecular insight, mutational analysis of hLF binding to hexon was carried out. Guided by the 2D class averages and the generated hybrid model, surface exposed and solvent accessible hLF residues in the regions of 2 apparent contact points were mutated, and binding affinities to fluorescently labelled hexon were measured using microscale thermophoresis (MST) (Fig. 4A). Single, charge-reducing mutations in Lfcin (mut1-4; R24S, R27A, K28A, and R30A) all resulted in reduced binding affinity of the respective hLF mutant. The importance of each of these residues was shown by their additive effect in a quadruple mutant (mut5; R24S/R27A/K28A/R30A) which showed the lowest binding affinity within the set of mutants, retaining less than 20% affinity of the WT protein. Point mutations were also introduced in a region directly adjacent to Lfcin which also harbors crosslink 1 (Hexon S446-hLF K282). Interestingly, this region, which has not been implicated in hexon binding before, also seems to play an important role for complex stability (Fig. 4A and B). A triple mutant (mut6; K277S/K280A/K285S) of hLF was found to be similarly impaired in its binding to hexon. Although, not measured, it is plausible that a mutant combining mutations from mut5 and 6 might not interact with hexon anymore.

**Figure 4.**
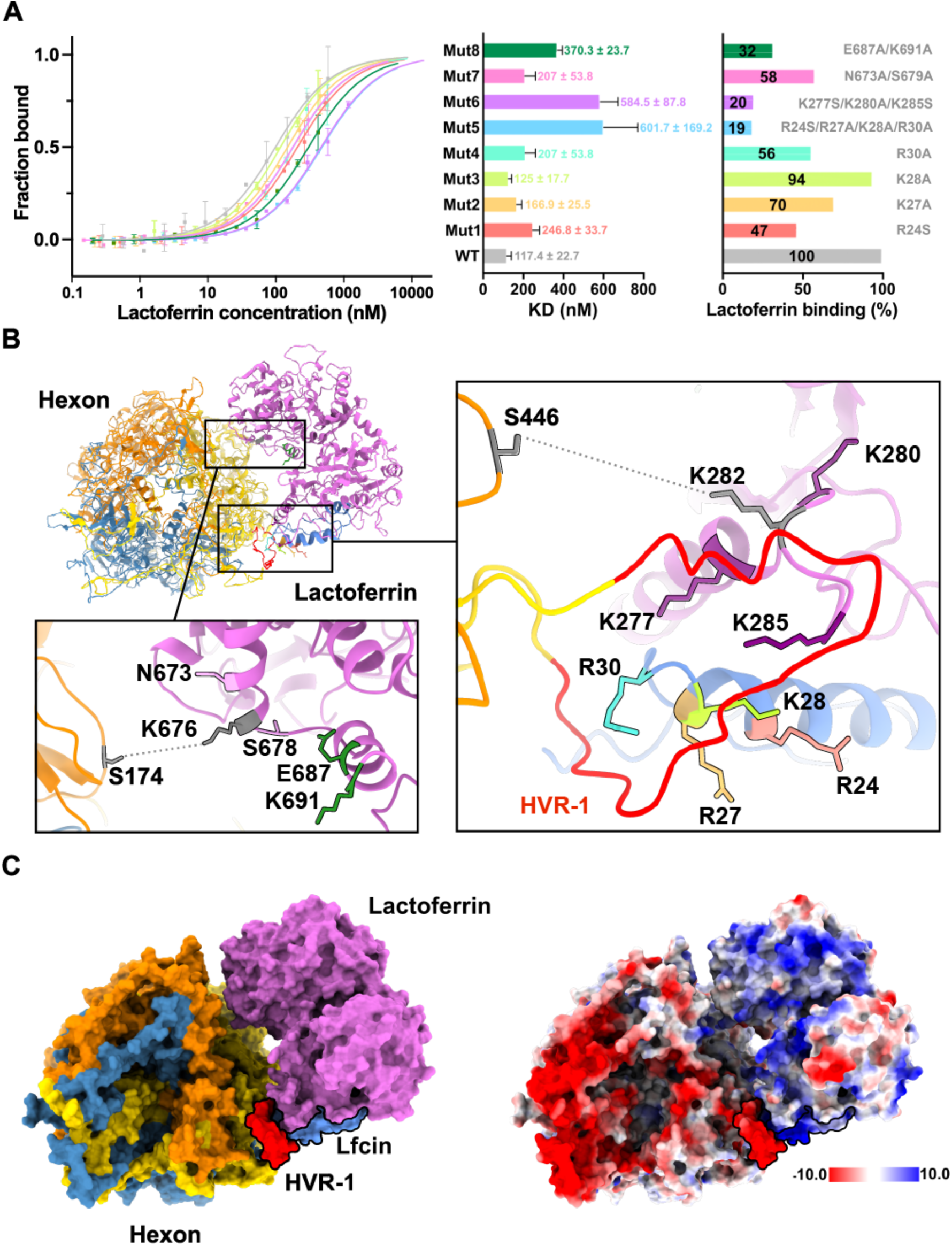
Validation of the hexon:hLF hybrid model. (A) Microscale Thermophoresis (MST) was used to determine the effect of mutations in hLF interface residues on the affinity to hexon. The fraction of hLF bound to fluorescently labelled hexon was determined and plotted as dose response curve (n=2). The binding curve of WT hLF is shown in grey, tested hLF mutants are shown in color. Colors of dose response curves are identical to colors used in bar graphs. The left diagram depicts the measured affinity; the right diagram shows the relative binding capacity of lactoferrin mutants to hexon in comparison to the WT. Percentage of lactoferrin binding is show inside the bar. Residues mutated in hLF are shown next to their respective bar. (B) Top view of the hexon:hLF hybrid model in cartoon representation. Mutants, with exception of Mut5, have been colored using the same color scheme as in (A). Additionally, HVR-1 on hexon has been colored red, Lfcin on lactoferrin is colored blue. Insets show close-up views of the mutated residues. Amino acids in identified cross-links are depicted in grey, cross-links are symbolically depicted as dotted lines. (C) Surface representation of the computed hexon:hLF complex in the colour scheme used throughout and colored according to the electrostatic potential. HVR-1 and LFcin have been highlighted on both representations with a black outline.

Double mutants mut7 (N673A/S678A) and mut8 (E687A/K691A) were introduced near the site of crosslink 2 which appears to be in proximity to hexon in our hybrid model (Fig. 4B and C) as well as 2D class averages (Fig. 3C). Although, to a lesser extent than mutations in the tip of the N-lobe, both double mutants are also significantly impaired in their binding to hexon. Binding affinities of these mutants are reduced by 40% and 70% for mut7 and mut8, respectively, pointing to a considerable role of the C-lobe in hexon binding (Fig. 4A).

## DISCUSSION

The human lung has developed an intricate system for protecting its epithelial lining against invading pathogens (33). For species C HAdVs the main mechanism for attachment and entry in respiratory epithelial cells uses CAR followed by secondary attachment to integrins that trigger uptake. This mechanism is well established and is likely the main port of infection in various immortalized cell lines but the relevance for the respiratory epithelium is questionable, given the hidden location of CAR deep within cellular junctions sealed off by tight junctions (19-21). Therefore, it is not surprising that alternative mechanisms for initial infections have been identified, such as using soluble components of various body fluids to gain cell entry. One such mechanism is to utilize hLF, and Lfcin, both of which are present in high concentrations within mucosal tissue (34). Naturally hLF is a protein of the innate protection of the lung and has been shown to interfere with infection ability of, for example, RSV (35) and HIV (36). In an inflamed tissue, neutrophils have been shown to secrete hLF, which apart from its role as an antimicrobial is also regarded as suppressor of inflammation (37). Therefore, it is very interesting that a human pathogen has evolved to highjack this system for infection. Unfortunately, very limited detailed information is available for this mechanism. In this study, we provide structural and biochemical data for the interaction between hLF and the HAdV-C5 hexon, which fill an important gap in our understanding of how HAdVs engage hLF for a CAR-independent infection of epithelial cells.

Previous studies have shown that the interaction with hLF likely involves the HVR-1 of hexon, which is the longest HVR of the protein (28). In addition, all species C adenoviruses have a longer HVR-1 containing multiple negatively charged residues. It has therefore been suggested that the length and charge of HVR-1 is essential for the ability to utilize lactoferrin for infection. We show here that the interaction between hLF and hexon is highly dependent on the ionic strength as buffers (20 mM HEPES) with NaCl concentrations above 5 mM disrupt the complex making it impossible to purify it by size exclusion chromatography. This was further verified by native mass spectrometry where a stable complex is present at ammonium acetate of 5-6 mM but not at 20 mM. These observations are somewhat puzzling from a biological context and challenge the biological relevance of the interaction. However, in both the lungs and the eye, tissues targeted by species C adenoviruses, there are local microenvironments present in which the ionic strength is regulated (38). The epithelial cell layer of the respiratory tract, including the trachea and bronchi, is covered by a thin layer of fluid, or mucus that ensures the proper function of the beating ciliated cells as well as protecting the epithelial cells from invading pathogens (39, 40). In this air-liquid interface the balance of ions such as sodium, chloride and bicarbonate is essential to preserve the characteristics of the mucins to ensure a normal mucociliary clearance (41, 42). Like in the conducting airways, the ionic strength in the tear film covering the cornea is tightly regulated to ensure proper function (43). Thus, as ions are pumped in and out of the cells, the local ionic strength is altered to a degree that may, at times, be beneficial for the hLF-mediated infection of the epithelium. In addition, very small differences in the ionic microenvironment may have large implications on the hLF-mediated entry given the avidity effect of the virions through the 240 copies of hexon trimers on each capsid (44). Even if the concentration of sodium has been determined in the sputum to be about 100 nM (+/- 27 nM) the concentration in the periciliary fluid is unknown and virtually impossible to determine (39).

The trimeric structure of HAdV-C5 hexon suggests a potential interaction with an equivalent number of hLF molecules in a 3:3 stoichiometric ratio to maximize avidity. Our native MS data, in agreement with 2D class averages of the complex, however, suggests a 3:1 stoichiometry of hexon to lactoferrin within the complex. Occasionally, a second molecule of lactoferrin was found to be bound, both in native MS (Fig. 1D) as well as in 2D projections (Fig. S6) while a 3:3 stoichiometry was not observed. A possible explanation could be an allosteric effect of the first binding event that introduces conformational changes in hexon resulting in reduced affinities for the second and third molecule of hLF. Alternatively, with 2 protomers of the hexon trimer already being occupied, binding of a third hLF might be sterically impaired.

An increasing amount of hybrid protein structures are currently being determined making use of an integrative approach combing experimental data and computational techniques (45). The structure of HAdV-C5 hexon in complex with human lactoferrin was well suited for this approach. Although we were unable to reconstruct a 3D model from our cryo-EM data alone, 2D class averages clearly showed assembly of the complex and identity of its components. Combining crosslinking data and refinement of the docked complex against the experimental 2D projections resulted in a likely model of the interaction. We believe that the workflow described here could also serve as a template to generate hybrid models of other difficult to obtain structures. While our hybrid model realistically depicts the overall assembly of the complex, it insufficiently describes direct contacts between interface residues in both structures. This is due to the inherent low resolution of the 2D class averages which only allow for rigid body refinement in addition to the high flexibility of hexon’s hypervariable loops in the hexon. These are typically largely disordered and therefore cannot be modelled or computationally predicted with accuracy. Furthermore, our hybrid model does not sufficiently take into account conformational changes that occur in both molecules upon binding. These result in small, observable disagreements between experimentally obtained 2D class averages and calculated 2D projections of the complex.

Further molecular insight was therefore obtained through site-directed mutagenesis of selected hLF residues present at its interface with hexon. Hereby we could confirm that a stretch of basic residues of hLF between amino acid positions 24-30 within the Lfcin region shows the highest contribution to the overall binding affinity. These residues appear to face the HVR-1 of hexon in the proposed model. This finding is in agreement with the earlier observation that an LFcin peptide comprising residues 1-18 did not increase HAdV-C5 transduction (28). Likewise, bovine LF (bLF), which carries substitutions in the mutated human lactoferrin residues, lacked transduction enhancing activity (28) (Fig. S7).

The described region forms a continuum with an adjacent patch of basic residues outside the Lfcin region which also contribute significantly to the binding affinity, or the lack thereof when mutated to Ala or Ser. It therefore seems that the region in the N-lobe of hLF that interacts with hexon HVR-1 might not be limited to the Lfcin peptide but comprises a larger area.

Interestingly, a possible second attachment point between hLF and hexon was observed in 2D class averages which appeared to involve the region around the junction of N- and C-lobe of hLF. The presence of such a site was corroborated by the identification of a cross-link connecting hLF K676 (present at the junction of N- and C-lobe) with hexon S174 (present in the ‘turrets’). Based on the electrostatic potential distribution in both molecules the interaction between the C-lobe of hLF and a second hexon protomer seems to be less charge dependent (Fig. 4E). Mutational analysis also indicated a significant, but lesser contribution to overall affinity within the complex. It is therefore possible that this second contact site is less crucial for overall complex stability, but rather for proper orientation of LF towards its cellular receptor.

Taken together our data provides novel structural insight into the interaction between HAdV-C5 and hLF. Combining computational modelling with cryo-EM, XL-MS and biophysical characterization, our hybrid model agrees well with mutational data, realistically depicting the interaction between HAdV-5C hexon and hLf. In the future, our results may be used for the generation of Lfcin-based antivirals that block the interaction between HVR-1 and hLF. In addition, novel Lfcin-based adjuvants may be developed that improve transduction efficiency in the lung during gene therapy.

## MATERIALS AND METHODS

### Cloning and protein purification

#### Human lactoferrin (hLF)

A commercially synthesised DNA fragment (Genewiz) encoding hLF (UniProt: Q19KS2) was cloned into the pHLsec vector (46) using the Gibson assembly method (New England Biolabs). Eight hLF mutants were produced by introducing desired point mutations during synthesis of DNA fragments (GenScript). The plasmids encoding the wild type (WT) LF, and its mutants were transfected and transiently expressed in Expi293F cells (ThermoFisher Scientific) following the manufacturer’s protocol.

LF and its mutants were purified from the culture supernatant obtained after centrifugation. After dialysis against 20 mM HEPES pH 7.2, 30 mM NaCl (buffer A) the supernatant was loaded onto a Mono S cation exchange chromatography column (GE Healthcare) equilibrated with buffer A using a ÄKTA Pure system (GE Healthcare). LF and its mutants were eluted by a linear NaCl gradient using buffer A and 20 mM HEPES pH 7.2, 1 M NaCl (buffer B) and subsequently analyzed by SDS-PAGE. Desired fractions were incubated with 0.5 mM FeCl_3_ in presence of 200 mM NaHCO_3_ for 1h at room temperature and further purified SEC on a S200 16/600 column (GE Healthcare) equilibrated with 20 mM HEPES pH 7.2, 150 mM NaCl (buffer C). The purified LF was concentrated using Amicon Ultra centrifugal filters (Merck Millipore) and stored at -80° C until further use.

#### Hexon

Hexon was purified from virus particles propagated in A549 cells (ATCC CCL-185). A549 cells were maintained in DMEM (Sigma) containing 5% (v/v) Fetal Bovine Serum (FBS; Gibco) and 1% (v/v) penicillin-streptomycin mix (Gibco) at 37 ° C under 5% CO_2_. Cells from a confluent culture were seeded into 5 x 175 cm^2^ flasks. Upon reaching confluence, medium was removed and fresh 5 ml medium containing 2% (v/v) FBS was added before infecting with HAdV-5C (200 µl). The flasks were incubated at 37 ° C for 90 min and gently mixed every 15 min. Later, the medium was replaced with fresh 30 ml medium containing 2% (v/v) Fetal Bovine Serum and flasks were left undisturbed for 4 days. Cells were harvested by tapping the flasks on the bench followed by centrifugation of the culture medium at 5,000 *g* for 10 min. Prior to centrifugation 1 ml culture from a flask was collected to be used as infectious material for a next round of hexon purification. A cell pellet from 5 x 175 cm^2^ was re-suspended in 5 ml 20 mM HEPES, 5 mM NaCl pH 7.2 (buffer D). To disrupt the cells, 3 freeze-thaw (37° C /-80° C) cycles were done and then 6 ml of Vertrel XF was added (1,1,1,2,3,4,4,5,5,5-decafluoropentane; SigmaAldrich). The cell suspension was mixed by gentle shaking of tubes by hands for 3 min followed by centrifugation at 3000 *g* for 10 min. The contents of clear top layer above the Vertrel phase were collected and separated over a CsCl gradient ultracentrifuged using SW41 rotor (Beckman Coulter) at 25000 rpm, 4°C for 90 min. The clear top phase obtained above a white band was collected and separated over a 5 ml HiTrap Q FF column (GE Healthcare) equilibrated with buffer A. Hexon was eluted with a linear NaCl gradient using buffers A and B. The elution fractions were analyzed by SDS-PAGE, concentrated and further purified by SEC on S200 16/600 column (GE Healthcare) equilibrated with buffer C. The purified hexon was stored at -80° C until further use.

#### Purification of hexon:hLF complex

Both hexon and hLF were dialyzed against buffer D before preparation of the complex. A 2-fold molar excess of LF was mixed with 925 pmoles of hexon and incubated on ice for 5 min followed by SEC on Superose 6 increase 10/300 (GE Healthcare) equilibrated with buffer D. Peak fractions were analyzed by SDS-PAGE to confirm the co-elution of hexon and hLF. The fractions containing the complex were concentrated to 1 mg/mL using a Vivaspin centrifugal concentrator (Sartorius).

### Loop prediction using AlphaFold

The structure of hexon was predicted using AlphaFold2 (47). Using PyMol (The PyMOL Molecular Graphics System, Version 2.0 Schrödinger, LLC.) and Coot (48) the loop regions 137-164, 188-190, 251-256, 271-278, and 431-436 from the hexon AlphaFold2 model were appended to the cryo-EM model lacking these regions.

### Native MS

Prior to native MS measurements, hexon:hLf complex purified in buffer D was buffer exchanged into 20 mM aqueous ammonium acetate solution pH 7.0 (AA; ≥99.99% trace metals basis, Honeywell Research Chemicals) using two cycles of spin gel-filtration (MicroBioSpin P-6, Bio-Rad). The complex was subsequently diluted by 20 mM AA to 0.5 µM (estimated based on 3+3 stoichiometry). Alternatively, for low ionic strength conditions, the dilution was performed with LC-MS grade water instead, resulting in 5.85 mM AA concentration. Reference spectra of isolated hexon and hLF were obtained from 5 µM (monomer) samples in 150 mM AA pH 7.0.

After short equilibration on ice, the samples were introduced into an Orbitrap Q Exactive UHMR mass spectrometer (Thermo Scientific) via static nanoelectrospray ionization. The samples were electrosprayed from gold-coated borosilicate glass capillaries Kwik-Fil 1B120F-4 (World Precision Instruments) prepared in-house essentially as described before (49, 50). The mass spectrometer was tuned for best signal quality and intensity, while maintaining minimal ion activation. Specifically, electrospray voltage was kept at 1.25 kV, source desolvation temperature 120°C, in-source trapping -50 V, ion transfer profile “high m/z”, analyzer profile “low m/z”, analyzer target resolution 1500 acquiring in mass range 1000 – 20000 m/z, averaging 5 microscans. Further desolvation and collisional cooling in HCD cell was performed with nitrogen at relative gas pressure setting 8.0 with gentle collisional activation by 10 V HCD voltage gradient. Raw spectra were averaged over at least 50 scans in Thermo Xcalibur Qual Browser 4.2.47 (Thermo Scientific) and mass deconvoluted in UniDec 6.0.3 package (51).

### Cross-linking MS

Hexon:hLF complex was cross-linked using the homobifunctional cross-linker disuccinimidyl dibutyric urea (DSBU; CF Plus Chemicals). The complex was mixed with excess of DSBU at 1:500 molar ratio. The reaction mixture was then incubated at room temperature for 1h and quenched with Tris at 30x higher concentration than DSBU.

Samples were diluted with 50 µL of 50 mM ammonium bicarbonate. Proteins were reduced by 10 mM TCEP and alkylated by 30 mM iodoacetamide. For deglycosylation of hLF, PNGase F (New England Biolabs) was added to the sample in 1:20 (w/w) ratio and incubated for 4h at 37°C. Proteins were digested by trypsin (Promega) over night at 37°C. After the digestion, samples were off line desalted using a Microtrap C18 cartridge (Optimize Technologies) and dried using a SpeedVac concentrator. Samples were re-suspended in 40 µl of 2% (v/v) acetonitrile, 0.1% (v/v) TFA and analysed using the Vanquish liquid chromatography system (ThermoScientific), connected to the timsToF SCP mass spectrometer equipped with captive spray (Bruker Daltonics). The mass spectrometer was operated in a positive data-dependent mode. 1 µl of the peptide mixture was injected by autosampler on the C18 trap column (PepMap Neo C18 5µm, 300 µm x 5 mm, Thermo Scientific). After 3 min of trapping, peptides were eluted from the trap column and separated on a C18 column (DNV PepMap Neo 75 µm x 150 mm, 2 µm, Thermo Scientific) by a linear 35 min water-acetonitrile gradient from 5 % (v/v) to 35 % (v/v) acetonitrile at a flow rate of 350 nl/min. The trap and analytical columns were both heated to 50°C. The TIMS scan range was set between 0.6 and 1.6 V s/ cm^2^ with a ramp time of 100 ms. The number of PASEF MS/MS ramps was 4. Precursor ions in the m/z range between 100 and 1700 with charge states ≥1+ and ≤8+ were selected for fragmentation. The active exclusion was enabled for 0.4 min.

The raw data were processed by Data Analysis 5.3 (Bruker Daltonics). The mgf files were uploaded to MeroX 2.0.1.4 software (52). The search parameters were set as follows: enzyme – trypsin (specific), carbamidomethylation as a fixed modification, oxidation of methionine as variable modifications. Precursor precision was set at 10.0 ppm and fragment ion precision was set at 0.1 Da.

### Cryo-EM sample preparation and data collection

#### HAdV-C5 hexon

3.5 μl of Hexon (0.7 mg/ml) was applied to freshly glow discharged Au, 300 mesh, R1.2/1.3 TEM grids (Protochips) coated in-house with a graphene monolayer. The grids were blotted at 100% humidity and 4° C (Vitrobot, ThemoFisher Scientific) and vitrified by plunging into liquid ethane. The grids were imaged using a Titan Krios G3i microscope (ThermoFisher Scientific) operated at 300 kV. Automated data collection was performed using EPU software (ThermoFisher Scientific) software. 1063 movies (40 frames /movie; 5 s exposure) were collected using a Quantum K2 LS detector (Gatan) at a magnification of 165.000x, resulting in a pixel size of 0.828 Å. The nominal defocus during data collection ranged from 1.2 µm - 2.5 µm and the total dose used was 55 e^−^/Å^2^.

#### HAdV-C5 hexon:hLF

Grids for cryo-EM were prepared by applying a freshly purified batch of hexon:hLF complex to Au, 300 mesh, R1.2/1.3 TEM grids (Quantifoil). 3 μl of Hexon-hLF complex (1 mg/ml) was applied to freshly glow discharged grids that were blotted at 100% humidity and 4°C (Vitrobot, ThemoFisher Scientific) and vitrified by plunging into liquid ethane. For imaging, the grids were transferred to a Talos Arctica microscope (ThermoFisher Scientific) equipped with a Falcon 3 detector (ThermoFisher Scientific). In total, 680 movies (60 frames/movies) were automatically collected using EPU software (ThermoFisher Scientific). The resulting pixel size was 1.23 Å and the nominal defocus ranged from 2 µm – 3.2 µm.

### Cryo-EM image processing

#### HAdV-C5 hexon

The recorded movies were entirely processed using cryoSPARC (version 3.2 and later) (53). Motion correction to obtain averaged micrographs from multi-frame movies followed by CTF estimation was performed using Patch Motion Correction, and CTFFIND4 algorithms integrated in cryoSPARC (53, 54). Hexon particles were picked using the Gaussian Blob Picker in cryoSPARC. Particles were extracted with a box size of 300 px and subjected to iterative, reference free 2D classification until no ill-defined class averages were obtained. The final selected subset of 98640 particles was used to produce 3 *ab initio* models. The class containing the maximum number of particles (53996) closely resembled the hexon and was chosen for homogeneous refinement with C3 symmetry imposed to yield the reconstructed hexon.

#### HAdV-C5 hexon:hLF

The recorded movies were imported into cryoSPARC (version 3.2 and later) and averaged micrographs from multi-frame movies were obtained via the Patch Motion Correction algorithm. Micrographs were CTF corrected using the Patch CTF Estimation algorithm. Blob Picker was employed to pick particles which were extracted with a box size of 230 px. The set of extracted particles was cleared of ill-defined particles by iterative, reference free 2D classification to obtain the final set of class averages, used for refinement of the HAdV-C5 hexon:hLF hybrid structure.

#### Model building HAdV-C5 hexon cryo-EM structure

To generate a model for hexon using the cryo-EM reconstruction, the crystal structure of hexon (PDB: 3TG7) was docked into the cryo-EM coloumb potential map using the dock_in_map procedure of Phenix (55). This docked model was subjected to automatic real-space refinement by Phenix. The obtained model was iteratively refined using multiple rounds of manual refinement performed in Coot followed by another round of automatic refinement in Phenix. After each round of refinement the model was validated using MolProbity (56) until satisfactory parameters were achieved.

### Molecular dynamics simulation HAdV-C5 hexon structure

#### System preparation

The model of hexon was trimmed down for more efficient calculation by discarding the rigid base, *i.e*. only residues 129-315 and 415-460 of all three protomers were retained in the final model. Hydrogens were added using the tLEaP program of AMBER20 suite (57). Titratable residues were modeled in their standard state at pH = 7.4. Protein force field ff19SB (58) was used along with a truncated octahedral box of SPC/E water molecules (59) extending 12 Å from the solute. Sodium and chloride counter ions were added. Prior to molecular dynamics (MD) simulation, the systems were relaxed, heated to 300 K and equilibrated at 300 K and 1 atm pressure using the published protocol (60).

#### Molecular dynamics

The production run was 600 ns long in NpT ensemble, i.e. keeping the number of particles (N), pressure (p) and temperature (T) constant. The SHAKE algorithm (61) was used to restrain all bond vibrations and hydrogen mass repartitioning to 3 Da (62) allowed us to apply a time step of 4 fs. Residues adjacent to those of the discarded base were fixed with harmonic restraint of 500 kcal mol^-1^ Å^-2^ to maintain the conformations compatible with linking to the base. All the analyses were done in the CPPTRAJ program (63).

### Docking of HAdV-C5 hexon:hLF complex and refinement of the docked model against 2D class averages

All simulations were performed in XPLOR-NIH v 3.4 (32) starting from the MD structure of Hexon and structure of hLF (PDB: 1LFG). Protons and missing side chain atoms in both proteins were added in XPLOR-NIH, followed by minimization of the energy function consisting of the standard geometric (bonds, angles, dihedrals, and impropers) and steric (van der Waals) terms. Based on the published analysis (64), the XL-MS data were converted into pairwise S446 (Hexon) – K282 (hLF) and S174 (Hexon) – K676 (hLF) intermolecular distance restraints, defined as the upper limit bounds of 24 Å between the Cα atoms and 15 Å bounds between the corresponding Ser OG and Lys NZ side chain atoms. The cryo-EM 2D class averages imported into XPLOR-NIH as two-dimensional coloumb potential maps (31, 65) were subsequently used to fit the protein molecules.

Following the computational protocol of Clore and co-workers (31), the refinement of the hexon:hLF complex was carried out in three steps. First, we performed a rigid-body protein-protein docking, driven by the intermolecular distance restraints. To this end, the positions of the hexon atoms were kept fixed, except for the “turret” loops (residues 136-164, 173-217, 268-287, and 429-448 in each subunit of the hexon homo-trimer, which includes the cross-link residues S174 and S446), which were given full torsional degrees of freedom. The incoming hLF molecule was treated as a rigid-body group, except for the cross-link residues K282 and K676, whose side chains were allowed to move. Starting from randomly oriented proteins separated by 100 Å, the docking comprised an initial rigid-body simulated annealing step, followed by the full side-chain energy minimization (32). In addition to the energy terms mentioned above, the total energy function included the intermolecular restraint term and a knowledge-based dihedral angle potential (32). In this first step, 100 structures were calculated and the 10 lowest-energy solutions – showing no distance restraint violations, imposed by crosslinks – retained for the subsequent refinement.

In the second step, energy terms for the 2D projections were added sequentially, with the map centering and (initially random) orientation optimized for each individual image from the cryo-EM 2D class stack. The back-calculated projections were computed as a sum over all heavy atoms, where each atom contributed a 2D quartic density according to its van der Waals radius and weighted by the atom’s number of electrons (31). The Gaussian blurring was employed throughout, with Gaussian diameter given by the resolution of the 2D class, and the agreement between the experimental and back-calculated electron density projections was estimated by the cross-correlation C metric (31). Out of 10 slices in the 2D classes image stack, one consistently failed to yield a good fit (C<0.9) and, thus, was discarded in the subsequent refinement rounds.

In the third and final step, the hexon:hLF complex was refined by allowing full rigid-body and torsional degrees of freedom in the motional setup described above, with simultaneous minimization of all constituent energy terms, including those for the intermolecular restraints and the nine individual electron density projections (31). In this run, 100 structures were calculated and 10 lowest-energy solutions – showing the best agreement with the experimental cryo-EM data (highest C values) and no distance restraint violations – retained for the final analysis. Individual back-calculated 2D projection images and their overlays with the experimental 2D class densities were output for visualization.

### Microscale thermophoresis

Microscale thermophoresis (MST) analysis was carried out to determine the binding affinity between hexon and hLF using Monolith NT.115 (Nanotemper Technologies). Hexon was labelled with AlexaFluor594 (ThermoFisher Scientific) according to manufacturer’s instructions. Excess dye was removed by SEC on S200 16/600 column (GE Healthcare) equilibrated with buffer C. MST experiments were performed in buffer D containing 0.05% (v/v) Tween 20. A range of 16 different concentrations of hLF or hLF mutants was generated by serial dilution (1:1) while the same amount of labelled hexon was added to each dilution. Following 5 min incubation at 25° C the samples were taken up in capillaries for MST measurements that were performed at 25° C at 50 % LED power, and 40 % MST power. The MST curves were fitted using MO.Affinity analysis software (Nanotemper Technologies).

## Data availability statement

The model of HAdV-C5 hexon has been deposited in the Protein Data Bank under accession code 8Q7C. The associated coulomb potential map has been deposited in the Electron Microscopy Data Bank under accession code EMD-18212.

The mass spectrometry proteomics data have been deposited to the ProteomeXchange Consortium via the PRIDE (66) partner repository with the dataset identifier PXD045900. Thirty snapshots from the MD trajectory used in integrative modeling are available at the Mendeley repository (doi: 10.17632/2xrxbm8g6j.1).

## Author contributions

NA and SZ conceptualized the study. AD carried out protein purifications and biophysical analyses. Cryo-EM data processing, subsequent model building, and molecular modelling was done by AD with input from HS and SK. OV performed the XPLOR-NIH analysis. AK carried out native MS measurements with input from CU. PP carried out XL-MS measurements. HS and AD prepared figures. SK prepared cryo-EM grids. ML performed MD simulations. KD supplied purified hexon and provided the purification protocol. SZ and BDP wrote the manuscript with input from all authors.

## Acknowledgements

We acknowledge Cryo-electron microscopy and tomography core facility CEITEC MU of CIISB, Instruct-CZ Centre, supported by MEYS CR (LM2023042) and European Regional Development Fund-Project „UP CIISB“(No. CZ.02.1.01/0.0/0.0/18_046/0015974) and iNEXT-Discovery, project number 871037, funded by the Horizon 2020 program of the European Commission.

We acknowledge Structural Mass Spectrometry core facility of CIISB, Instruct-CZ Centre, supported by MEYS CR (LM2023042) and European Regional Development Fund-Project „UP CIISB“ (No. CZ.02.1.01/0.0/0.0/18_046/0015974). HS was supported by the Grant Agency of Charles University (project no. 383821/2600). AD was supported by the IOCB Postdoctoral Fellowship. CU acknowledges support through EU Horizon 2020 ERC StG-2017 759661.

## References

1. Greber UF, Flatt JW. 2019. Adenovirus Entry: From Infection to Immunity. Annu Rev Virol 6:177–197.

2. Benko M, Aoki K, Arnberg N, Davison AJ, Echavarria M, Hess M, Jones MS, Kajan GL, Kajon AE, Mittal SK, Podgorski, II, San Martin C, Wadell G, Watanabe H, Harrach B, Ictv Report C. 2022. ICTV Virus Taxonomy Profile: Adenoviridae 2022. J Gen Virol 103.

3. Echavarria M. 2008. Adenoviruses in immunocompromised hosts. Clin Microbiol Rev 21:704–15.

4. Proenca-Modena JL, de Souza Cardoso R, Criado MF, Milanez GP, de Souza WM, Parise PL, Bertol JW, de Jesus BLS, Prates MCM, Silva ML, Buzatto GP, Demarco RC, Valera FCP, Tamashiro E, Anselmo-Lima WT, Arruda E. 2019. Human adenovirus replication and persistence in hypertrophic adenoids and palatine tonsils in children. J Med Virol 91:1250–1262.

5. Lion T. 2019. Adenovirus persistence, reactivation, and clinical management. FEBS Lett 593:3571–3582.

6. Ohnishi Y, Noda S. 1976. [Studies on the rabbit conjunctival cells in vitro (author’s transl)]. Nippon Ganka Gakkai Zasshi 80:1255–63.

7. Berk AJ, Sharp PA. 1978. Structure of the adenovirus 2 early mRNAs. Cell 14:695–711.

8. Cepko CL, Sharp PA. 1982. Assembly of adenovirus major capsid protein is mediated by a nonvirion protein. Cell 31:407–15.

9. Lee A. 2023. Nadofaragene Firadenovec: First Approval. Drugs 83:353–357.

10. Sumida SM, Truitt DM, Kishko MG, Arthur JC, Jackson SS, Gorgone DA, Lifton MA, Koudstaal W, Pau MG, Kostense S, Havenga MJ, Goudsmit J, Letvin NL, Barouch DH. 2004. Neutralizing antibodies and CD8+ T lymphocytes both contribute to immunity to adenovirus serotype 5 vaccine vectors. J Virol 78:2666–73.

11. Tillman BW, de Gruijl TD, Luykx-de Bakker SA, Scheper RJ, Pinedo HM, Curiel TJ, Gerritsen WR, Curiel DT. 1999. Maturation of dendritic cells accompanies high-efficiency gene transfer by a CD40-targeted adenoviral vector. J Immunol 162:6378–83.

12. Milligan ID, Gibani MM, Sewell R, Clutterbuck EA, Campbell D, Plested E, Nuthall E, Voysey M, Silva-Reyes L, McElrath MJ, De Rosa SC, Frahm N, Cohen KW, Shukarev G, Orzabal N, van Duijnhoven W, Truyers C, Bachmayer N, Splinter D, Samy N, Pau MG, Schuitemaker H, Luhn K, Callendret B, Van Hoof J, Douoguih M, Ewer K, Angus B, Pollard AJ, Snape MD. 2016. Safety and Immunogenicity of Novel Adenovirus Type 26- and Modified Vaccinia Ankara-Vectored Ebola Vaccines: A Randomized Clinical Trial. JAMA 315:1610–23.

13. Barouch DH, Tomaka FL, Wegmann F, Stieh DJ, Alter G, Robb ML, Michael NL, Peter L, Nkolola JP, Borducchi EN, Chandrashekar A, Jetton D, Stephenson KE, Li W, Korber B, Tomaras GD, Montefiori DC, Gray G, Frahm N, McElrath MJ, Baden L, Johnson J, Hutter J, Swann E, Karita E, Kibuuka H, Mpendo J, Garrett N, Mngadi K, Chinyenze K, Priddy F, Lazarus E, Laher F, Nitayapan S, Pitisuttithum P, Bart S, Campbell T, Feldman R, Lucksinger G, Borremans C, Callewaert K, Roten R, Sadoff J, Scheppler L, Weijtens M, Feddes-de Boer K, van Manen D, Vreugdenhil J, Zahn R, Lavreys L, et al. 2018. Evaluation of a mosaic HIV-1 vaccine in a multicentre, randomised, double-blind, placebo-controlled, phase 1/2a clinical trial (APPROACH) and in rhesus monkeys (NHP 13-19). Lancet 392:232–243.

14. Larocca RA, Mendes EA, Abbink P, Peterson RL, Martinot AJ, Iampietro MJ, Kang ZH, Aid M, Kirilova M, Jacob-Dolan C, Tostanoski L, Borducchi EN, De La Barrera RA, Barouch DH. 2019. Adenovirus Vector-Based Vaccines Confer Maternal-Fetal Protection against Zika Virus Challenge in Pregnant IFN-alphabetaR(-/-) Mice. Cell Host Microbe 26:591–600 e4.

15. Williams K, Bastian AR, Feldman RA, Omoruyi E, de Paepe E, Hendriks J, van Zeeburg H, Godeaux O, Langedijk JPM, Schuitemaker H, Sadoff J, Callendret B. 2020. Phase 1 Safety and Immunogenicity Study of a Respiratory Syncytial Virus Vaccine With an Adenovirus 26 Vector Encoding Prefusion F (Ad26.RSV.preF) in Adults Aged >/=60 Years. J Infect Dis 222:979-988.

16. Sadoff J, Gray G, Vandebosch A, Cardenas V, Shukarev G, Grinsztejn B, Goepfert PA, Truyers C, Fennema H, Spiessens B, Offergeld K, Scheper G, Taylor KL, Robb ML, Treanor J, Barouch DH, Stoddard J, Ryser MF, Marovich MA, Neuzil KM, Corey L, Cauwenberghs N, Tanner T, Hardt K, Ruiz-Guinazu J, Le Gars M, Schuitemaker H, Van Hoof J, Struyf F, Douoguih M, Group ES. 2021. Safety and Efficacy of Single-Dose Ad26.COV2.S Vaccine against Covid-19. N Engl J Med 384:2187-2201.

17. Bergelson JM, Cunningham JA, Droguett G, Kurt-Jones EA, Krithivas A, Hong JS, Horwitz MS, Crowell RL, Finberg RW. 1997. Isolation of a common receptor for Coxsackie B viruses and adenoviruses 2 and 5. Science 275:1320–3.

18. Storm RJ, Persson BD, Skalman LN, Frangsmyr L, Lindstrom M, Rankin G, Lundmark R, Domellof FP, Arnberg N. 2017. Human Adenovirus Type 37 Uses alpha(V)beta(1) and alpha(3)beta(1) Integrins for Infection of Human Corneal Cells. J Virol 91.

19. Mateo M, Generous A, Sinn PL, Cattaneo R. 2015. Connections matter--how viruses use cell-cell adhesion components. J Cell Sci 128:431–9.

20. Walters RW, Freimuth P, Moninger TO, Ganske I, Zabner J, Welsh MJ. 2002. Adenovirus fiber disrupts CAR-mediated intercellular adhesion allowing virus escape. Cell 110:789–99.

21. Walters RW, Grunst T, Bergelson JM, Finberg RW, Welsh MJ, Zabner J. 1999. Basolateral localization of fiber receptors limits adenovirus infection from the apical surface of airway epithelia. J Biol Chem 274:10219–26.

22. Kotha PL, Sharma P, Kolawole AO, Yan R, Alghamri MS, Brockman TL, Gomez-Cambronero J, Excoffon KJ. 2015. Adenovirus entry from the apical surface of polarized epithelia is facilitated by the host innate immune response. PLoS Pathog 11:e1004696.

23. Waddington SN, McVey JH, Bhella D, Parker AL, Barker K, Atoda H, Pink R, Buckley SM, Greig JA, Denby L, Custers J, Morita T, Francischetti IM, Monteiro RQ, Barouch DH, van Rooijen N, Napoli C, Havenga MJ, Nicklin SA, Baker AH. 2008. Adenovirus serotype 5 hexon mediates liver gene transfer. Cell 132:397–409.

24. Lyle C, McCormick F. 2010. Integrin alphavbeta5 is a primary receptor for adenovirus in CAR-negative cells. Virol J 7:148.

25. Cheneau C, Eichholz K, Tran TH, Tran TTP, Paris O, Henriquet C, Bajramovic JJ, Pugniere M, Kremer EJ. 2021. Lactoferrin Retargets Human Adenoviruses to TLR4 to Induce an Abortive NLRP3-Associated Pyroptotic Response in Human Phagocytes. Front Immunol 12:685218.

26. Bellamy W, Takase M, Yamauchi K, Wakabayashi H, Kawase K, Tomita M. 1992. Identification of the bactericidal domain of lactoferrin. Biochim Biophys Acta 1121:130–6.

27. Johansson C, Jonsson M, Marttila M, Persson D, Fan XL, Skog J, Frangsmyr L, Wadell G, Arnberg N. 2007. Adenoviruses use lactoferrin as a bridge for CAR-independent binding to and infection of epithelial cells. J Virol 81:954–63.

28. Persson BD, Lenman A, Frangsmyr L, Schmid M, Ahlm C, Pluckthun A, Jenssen H, Arnberg N. 2020. Lactoferrin-Hexon Interactions Mediate CAR-Independent Adenovirus Infection of Human Respiratory Cells. J Virol 94.

29. Jonsson MI, Lenman AE, Frangsmyr L, Nyberg C, Abdullahi M, Arnberg N. 2009. Coagulation factors IX and X enhance binding and infection of adenovirus types 5 and 31 in human epithelial cells. J Virol 83:3816–25.

30. Lenman A, Muller S, Nygren MI, Frangsmyr L, Stehle T, Arnberg N. 2011. Coagulation factor IX mediates serotype-specific binding of species A adenoviruses to host cells. J Virol 85:13420–31.

31. Walti MA, Canagarajah B, Schwieters CD, Clore GM. 2021. Visualization of Sparsely-populated Lower-order Oligomeric States of Human Mitochondrial Hsp60 by Cryo-electron Microscopy. J Mol Biol 433:167322.

32. Schwieters CD, Kuszewski JJ, Tjandra N, Clore GM. 2003. The Xplor-NIH NMR molecular structure determination package. J Magn Reson 160:65–73.

33. Martin TR, Frevert CW. 2005. Innate immunity in the lungs. Proc Am Thorac Soc 2:403–11.

34. Rosa L, Cutone A, Lepanto MS, Paesano R, Valenti P. 2017. Lactoferrin: A Natural Glycoprotein Involved in Iron and Inflammatory Homeostasis. Int J Mol Sci 18.

35. Sano H, Nagai K, Tsutsumi H, Kuroki Y. 2003. Lactoferrin and surfactant protein A exhibit distinct binding specificity to F protein and differently modulate respiratory syncytial virus infection. Eur J Immunol 33:2894–902.

36. Berlutti F, Pantanella F, Natalizi T, Frioni A, Paesano R, Polimeni A, Valenti P. 2011. Antiviral properties of lactoferrin--a natural immunity molecule. Molecules 16:6992–7018.

37. Lonnerdal B, Iyer S. 1995. Lactoferrin: molecular structure and biological function. Annu Rev Nutr 15:93–110.

38. Toczylowska-Maminska R, Dolowy K. 2012. Ion transporting proteins of human bronchial epithelium. J Cell Biochem 113:426–32.

39. Luk CK, Dulfano MJ. 1983. Effect of pH, viscosity and ionic-strength changes on ciliary beating frequency of human bronchial explants. Clin Sci (Lond) 64:449–51.

40. Tabary O, Muselet C, Miesch MC, Yvin JC, Clement A, Jacquot J. 2003. Reduction of chemokine IL-8 and RANTES expression in human bronchial epithelial cells by a sea-water derived saline through inhibited nuclear factor-kappaB activation. Biochem Biophys Res Commun 309:310–6.

41. Fajac I, Burgel PR. 2023. Cystic fibrosis. Presse Med 52:104169.

42. Saint-Criq V, Gray MA. 2017. Role of CFTR in epithelial physiology. Cell Mol Life Sci 74:93–115.

43. Hosoya K, Lee VH, Kim KJ. 2005. Roles of the conjunctiva in ocular drug delivery: a review of conjunctival transport mechanisms and their regulation. Eur J Pharm Biopharm 60:227–40.

44. Gallardo J, Perez-Illana M, Martin-Gonzalez N, San Martin C. 2021. Adenovirus Structure: What Is New? Int J Mol Sci 22.

45. Vallat B, Webb B, Fayazi M, Voinea S, Tangmunarunkit H, Ganesan SJ, Lawson CL, Westbrook JD, Kesselman C, Sali A, Berman HM. 2021. New system for archiving integrative structures. Acta Crystallogr D Struct Biol 77:1486–1496.

46. Aricescu AR, Lu W, Jones EY. 2006. A time- and cost-efficient system for high-level protein production in mammalian cells. Acta Crystallogr D Biol Crystallogr 62:1243–50.

47. Mirdita M, Schutze K, Moriwaki Y, Heo L, Ovchinnikov S, Steinegger M. 2022. ColabFold: making protein folding accessible to all. Nat Methods 19:679–682.

48. Emsley P, Lohkamp B, Scott WG, Cowtan K. 2010. Features and development of Coot. Acta Crystallogr D Biol Crystallogr 66:486–501.

49. Hernandez H, Robinson CV. 2007. Determining the stoichiometry and interactions of macromolecular assemblies from mass spectrometry. Nat Protoc 2:715–26.

50. Krichel B, Bylapudi G, Schmidt C, Blanchet C, Schubert R, Brings L, Koehler M, Zenobi R, Svergun D, Lorenzen K, Madhugiri R, Ziebuhr J, Uetrecht C. 2021. Hallmarks of Alpha- and Betacoronavirus non-structural protein 7+8 complexes. Sci Adv 7.

51. Marty MT, Baldwin AJ, Marklund EG, Hochberg GK, Benesch JL, Robinson CV. 2015. Bayesian deconvolution of mass and ion mobility spectra: from binary interactions to polydisperse ensembles. Anal Chem 87:4370–6.

52. Iacobucci C, Gotze M, Ihling CH, Piotrowski C, Arlt C, Schafer M, Hage C, Schmidt R, Sinz A. 2018. A cross-linking/mass spectrometry workflow based on MS-cleavable cross-linkers and the MeroX software for studying protein structures and protein-protein interactions. Nat Protoc 13:2864–2889.

53. Punjani A, Rubinstein JL, Fleet DJ, Brubaker MA. 2017. cryoSPARC: algorithms for rapid unsupervised cryo-EM structure determination. Nat Methods 14:290–296.

54. Rohou A, Grigorieff N. 2015. CTFFIND4: Fast and accurate defocus estimation from electron micrographs. J Struct Biol 192:216–21.

55. Liebschner D, Afonine PV, Baker ML, Bunkoczi G, Chen VB, Croll TI, Hintze B, Hung LW, Jain S, McCoy AJ, Moriarty NW, Oeffner RD, Poon BK, Prisant MG, Read RJ, Richardson JS, Richardson DC, Sammito MD, Sobolev OV, Stockwell DH, Terwilliger TC, Urzhumtsev AG, Videau LL, Williams CJ, Adams PD. 2019. Macromolecular structure determination using X-rays, neutrons and electrons: recent developments in Phenix. Acta Crystallogr D Struct Biol 75:861–877.

56. Chen VB, Arendall WB, 3rd, Headd JJ, Keedy DA, Immormino RM, Kapral GJ, Murray LW, Richardson JS, Richardson DC. 2010. MolProbity: all-atom structure validation for macromolecular crystallography. Acta Crystallogr D Biol Crystallogr 66:12-21.

57. D.A. Case HMA, K. Belfon, I.Y. Ben-Shalom, J.T. Berryman, S.R. Brozell, D.S. Cerutti, T.E. Cheatham, III, G.A. Cisneros, V.W.D. Cruzeiro, T.A. Darden, N. Forouzesh, G. Giambaşu, T. Giese, M.K. Gilson, H. Gohlke, A.W. Goetz, J. Harris, S. Izadi, S.A. Izmailov, K. Kasavajhala, M.C. Kaymak, E. King, A. Kovalenko, T. Kurtzman, T.S. Lee, P. Li, C. Lin, J. Liu, T. Luchko, R. Luo, M. Machado, V. Man, M. Manathunga, K.M. Merz, Y. Miao, O. Mikhailovskii, G. Monard, H. Nguyen, K.A. O’Hearn, A. Onufriev, F. Pan, S. Pantano, R. Qi, A. Rahnamoun, D.R. Roe, A. Roitberg, C. Sagui, S. Schott-Verdugo, A. Shajan, J. Shen, C.L. Simmerling, N.R. Skrynnikov, J. Smith, J. Swails, R.C. Walker, J. Wang, J. Wang, H. Wei, X. Wu, Y. Wu, Y. Xiong, Y. Xue, D.M. York, S. Zhao, Q. Zhu, and P.A. Kollman 2020. Amber 2020, University of California, San Francisco.

58. Tian C, Kasavajhala K, Belfon KAA, Raguette L, Huang H, Migues AN, Bickel J, Wang Y, Pincay J, Wu Q, Simmerling C. 2020. ff19SB: Amino-Acid-Specific Protein Backbone Parameters Trained against Quantum Mechanics Energy Surfaces in Solution. J Chem Theory Comput 16:528–552.

59. Berendsen HJCG, J. R.; Straatsma, T. P.. 1987. The missing term in effective pair potentials. The Journal of Physical Chemistry 91:6269–6271.

60. Srb P, Svoboda M, Benda L, Lepsik M, Tarabek J, Sicha V, Gruner B, Grantz-Saskova K, Brynda J, Rezacova P, Konvalinka J, Veverka V. 2019. Capturing a dynamically interacting inhibitor by paramagnetic NMR spectroscopy. Phys Chem Chem Phys 21:5661–5673.

61. Ryckaert J-P, Ciccotti G, Berendsen HJC. 1977. Numerical integration of the cartesian equations of motion of a system with constraints: molecular dynamics of n-alkanes. Journal of Computational Physics 23:327–341.

62. Hopkins CW, Le Grand S, Walker RC, Roitberg AE. 2015. Long-Time-Step Molecular Dynamics through Hydrogen Mass Repartitioning. J Chem Theory Comput 11:1864–74.

63. Roe DR, Cheatham TE, 3rd. 2013. PTRAJ and CPPTRAJ: Software for Processing and Analysis of Molecular Dynamics Trajectory Data. J Chem Theory Comput 9:3084-95.

64. Merkley ED, Rysavy S, Kahraman A, Hafen RP, Daggett V, Adkins JN. 2014. Distance restraints from crosslinking mass spectrometry: mining a molecular dynamics simulation database to evaluate lysine-lysine distances. Protein Sci 23:747–59.

65. Schwieters CD, Bermejo GA, Clore GM. 2018. Xplor-NIH for molecular structure determination from NMR and other data sources. Protein Sci 27:26–40.

66. Perez-Riverol Y, Bai J, Bandla C, Garcia-Seisdedos D, Hewapathirana S, Kamatchinathan S, Kundu DJ, Prakash A, Frericks-Zipper A, Eisenacher M, Walzer M, Wang S, Brazma A, Vizcaino JA. 2022. The PRIDE database resources in 2022: a hub for mass spectrometry-based proteomics evidences. Nucleic Acids Res 50:D543–D552.

