## Supplementary information for "Structural Insights into the Interaction Between Adenovirus C5 Hexon and Human Lactoferrin"

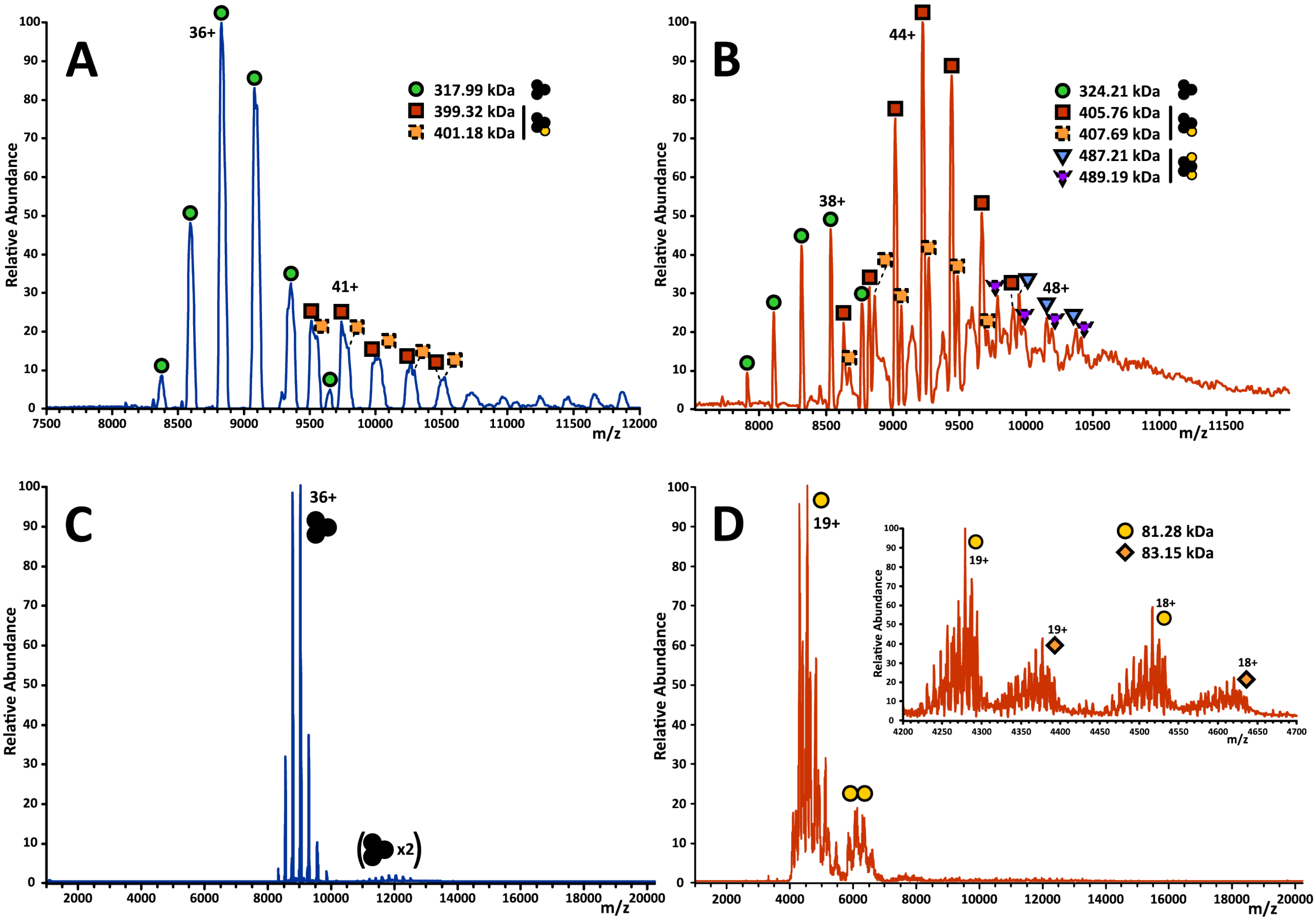


**Fig. S1.** Detailed native MS peak annotation. Close-up at higher (3.000) instrument resolving power setting shows details of detected 0.5 µM hexon:hLF complex species in 20 mM (A) and 5.85 mM (B) ammonium acetate pH 7.0. While hexon alone (C) is trimeric and highly homogeneous (5 µM in 150 mM ammonium acetate pH 7.0), hLF alone (D - at 5 µM in 150 mM AA pH 7.0) shows major heterogeneity at 12.000 MS resolving power setting stemming from the presence of two major proteoforms differing by 1.86 kDa as well as a broad distribution of hexose units (inset). This in turn explains the broad multi-apex peaks of hLF-containing complexes detected in A and B.

**Table S1.** Cryo-EM data collection, and model quality parameters

| | **HAdV-C5 hexon** (EMD- 18212) (PDB- 8Q7C) |
|---|---|
| **Data collection and processing** | |
| Microscope | Titan Krios |
| Detector | K2 LS |
| Magnification (nominal) | 165.000x |
| Voltage (kV) | 300 |
| Spherical abberation | 2.7 mm |
| Total electron dose ($e^{-}$/Å$^2$) | 55 |
| Defocus range (µm) | -2.5 to 1.2 |
| Pixel size (Å) | 0.82 |
| Number of Micrographs | 1063 |
| Final particle images (no.) | 53396 |
| Map resolution (Å)<br>FSC threshold | 2.96 ($FSC_{0.143}$) |
| **Refinement** | |
| Initial model used (PDB code) | 3TG7 |
| RMSZ | |
| Bond lengths | 0.31 |
| Bond angles | 0.50 |
| Validation | |
| MolProbity score | 2.03 |
| Clashscore, all-atom | 7.61 |
| Poor rotamers | 3.7% |
| Ramachandran plot | |
| Favored | 96.92% |
| Allowed | 3.08% |
| Outliers | 0% |

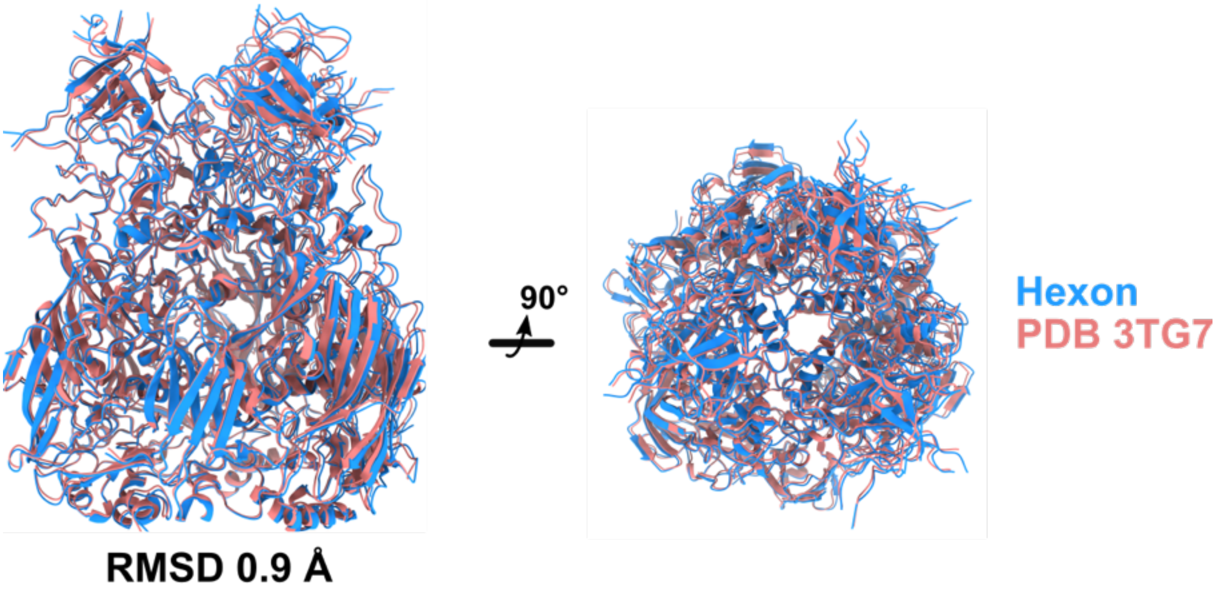


**Fig. S2.** Comparison of the HAdV-C5 hexon model obtained by single-particle cryoEM with a crystal structure of the same protein. The hexon model built using the cryo-EM reconstruction is depicted in blue (PDB: 8Q7C), the crystal structure (PDB: 3TG7) is shown in pink. Both structures are almost identical and can be aligned to an RMSD of 0.9Å.

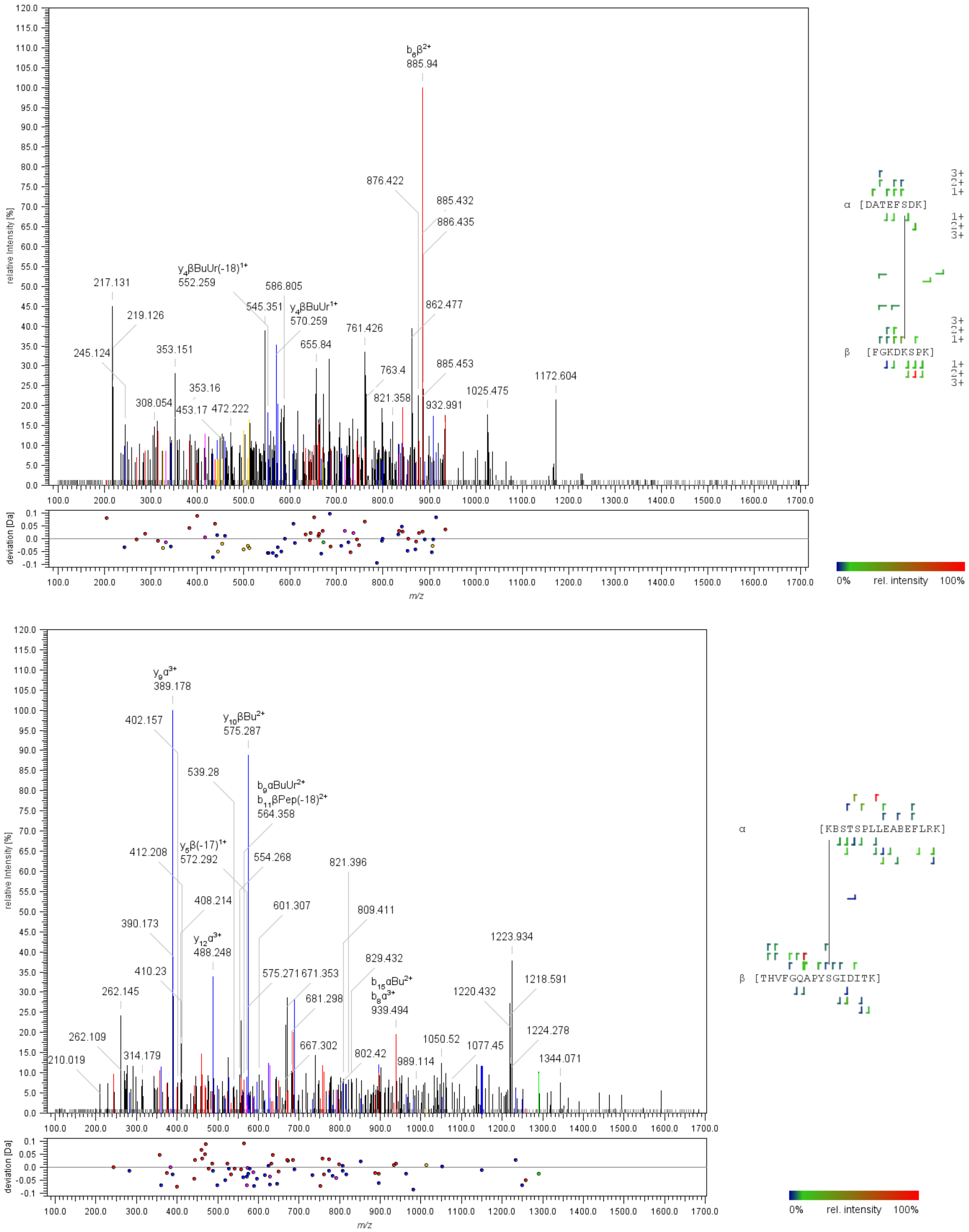


**Fig. S3.** Annotated tandem mass spectra of two intermolecular cross-linked peptides by MS-cleavable homobifunctional disuccinimidyl dibutyric urea (DBSU) reagent. Several peptide backbone and peptide-cross-linker (Bu and BuUr) fragments were matched using the MeroX software.

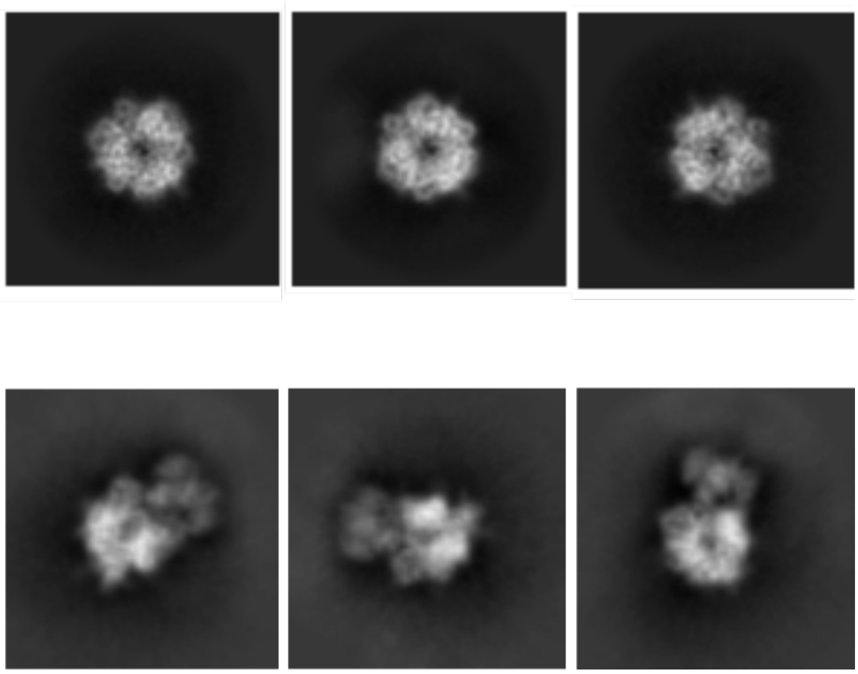

**Fig. S4.** 2D class averages showing hexon without no hLF bound (top), and bound to hLF (below). Conformational changes in hexon suggest an ‘opening-up’ of the hexon top upon binding of hLF.

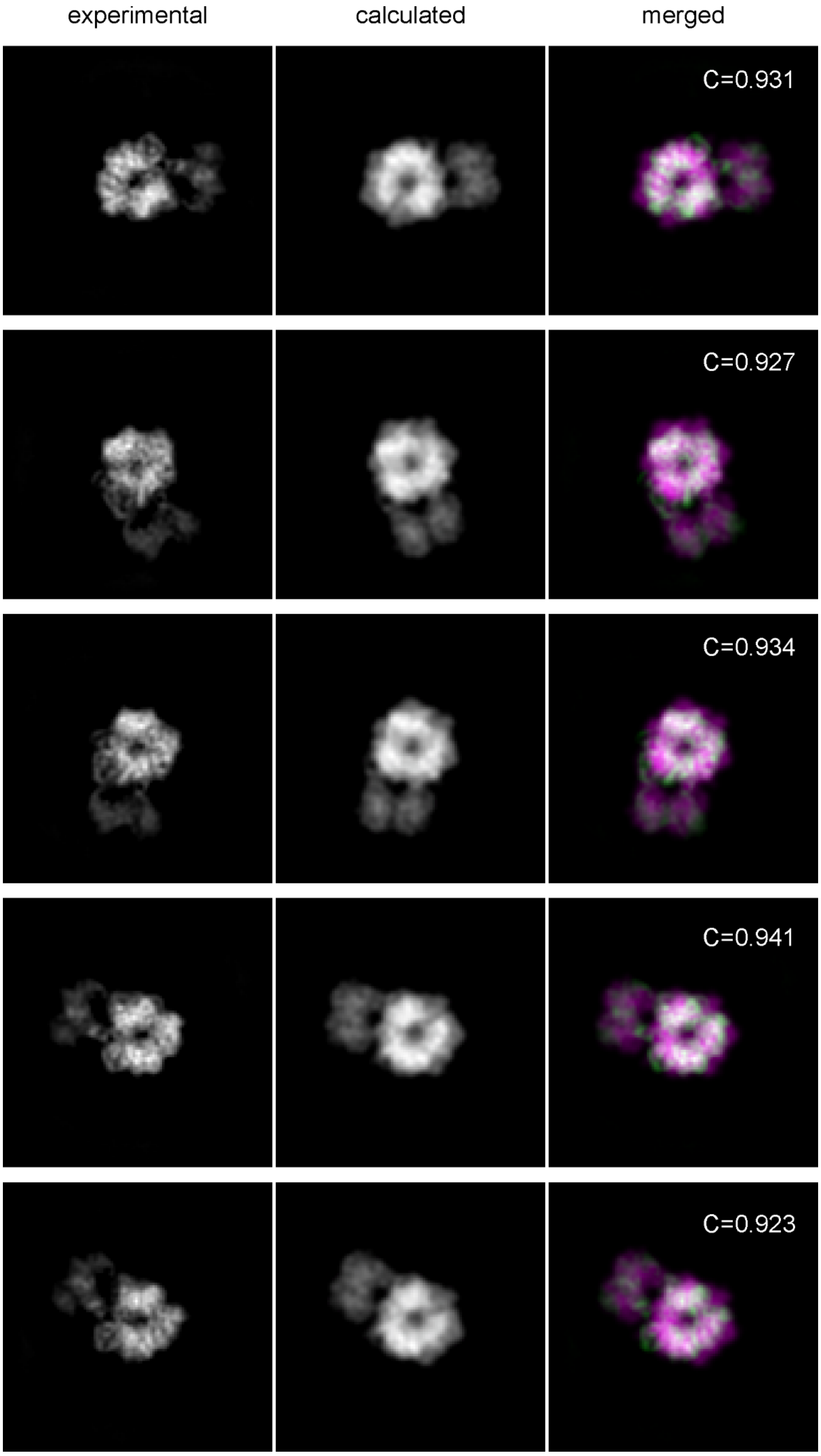


**Fig. S5.** Remaining lowest-energy solutions of hexon:hLF complex refined against 2D class averages of the complex. The left column shows the experimentally obtained 2D class averages for the hexon:hLF, the central column shows the calculated back projections for the hexon:hLF complex obtained after docking and energy minimization, and the right column shows the result of superposition of experimental (green) and calculated (magenta) projections. Cross-correlation (C) of experimental and calculated projections is shown for individual projections

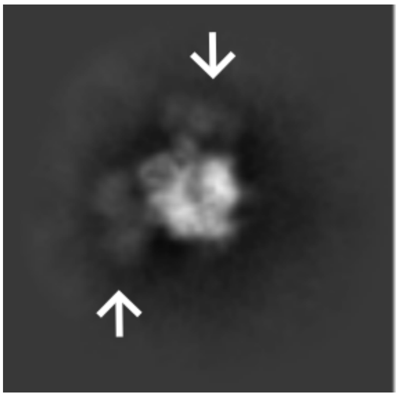

**Fig. S6.** A 2D class average showing 2 hLF copies (white arrows) bound to a hexon trimer.

```
hLF      GRRRSVQWCAVSQPEATKCFQWQRNMRKVRGPPVSCIKRDSPIQCIQAIAENRADAVTLD      60
bLF      APRKNVRWCTISQPEWFKCRRWQWRMKKLGAPSITCVRRAFALECIRAIAEKKADAVTLD      60
         . *:.*:**::****  ** :** .*:*: .* ::*::*   ::**:****::*******

hLF      GGFIYEAGLAPYKLRPVAAEVYGTERQPRTHYYAVAVVKKGGSFQLNELQGLKSCHTGLR      120
bLF      GGMVFEAGRDPYKLRPVAAEIYGTKESPQTHYYAVAVVKKGSNFQLDQLQGRKSCHTGLG      120
         **::***  **********:***:..*:************..***::*** *******

hLF      RTAGWNVPIGTLRPFLNWTGPPEPIEAAVARFFSASCVPGADKGQFPNLCRLCAGTGENK      180
bLF      RSAGWIIPMGILRPYLSWTESLEPLQGAVAKFFSASCVPCIDRQAYPNLCQLCKGEGENQ      180
         *:*** :*:* ***:*.**   **::.***:********  *:  :****:** * ***:

hLF      CAFSSQEPYFSYSGAFKCLRDGAGDVAFIRESTVFEDLSDEAERDEYELLCPDNTRKPVD      240
bLF      CACSSREPYFGYSGAFKCLQDGAGDVAFVKETTVFENLPEKADRDQYELLCLNNSRAPVD      240
         ** **:****.********:********::*:****:* ::*:**:***** :*:* ***

hLF      KFKDCHLARVPSHAVVARSVNGKEDAIWNLLRQAQEKFGKDKSPKFQLFGSPSGQKDLLF      300
bLF      AFKECHLAQVPSHAVVARSVDGKEDLIWKLLSKAQEKFGKNKSRSFQLFGSPPGQRDLLF      300
          **:****:***********:**** **:** :*******:** .******* **:****

hLF      KDSAIGFSRVPPRIDSGLYLGSGYFTAIQNLRKSEEEVAARRARVVWCAVGEQELRKCNQ      360
bLF      KDSALGFLRIPSKVDSALYLGSRYLTTLKNLRETAEEVKARYTRVVWCAVGPEEQKKCQQ      360
         ****:** *:* ::**.***** *:*::***:: *** ** :******** :* :**:*

hLF      WSGLSEGSVTCSSASTTEDCIALVLKGEADAMSLDGGYVYTAGKCGLVPVLAENYKSQQS      420
bLF      WSQQSGQNVTCATASTTDDCIVLVLKGEADALNLDGGYIYTAGKCGLVPVLAENRKSSKH      420
         **  *  .***::****:***.*********:.*****:*************** **.:

hLF      SDPDPNCVDRPVEGYLAVAVVRRSDTSLTWNSVKGKKSCHTAVDRTAGWNIPMGLLFNQT      480
bLF      --SSLDCVLRPTEGYLAVAVVKKANEGLTWNSLKDKKSCHTAVDRTAGWNIPMGLIVNQT      478
            . :** **.*********:::: .*****:*.********************:.***

hLF      GSCKFDEYFSQSCAPGSDPRSNLCALCIGDEQGENKCVPNSNERYYGYTGAFRCLAENAG      540
bLF      GSCAFDEFFSQSCAPGADPKSRLCALCAGDDQGLDKCVPNSKEKYYGYTGAFRCLAEDVG      538
         *** ***:********:**:*.***** **:** :******:*:*************:.*

hLF      DVAFVKDVTVLQNTDGNNNEAWAKDLKLADFALLCLDGKRKPVTEARSCHLAMAPNHAVV      600
bLF      DVAFVKNDTVWENTNGESTADWAKNLNREDFRLLCLDGTRKPVTEAQSCHLAVAPNHAVV      598
         ******: ** :**:*:..  ***:*:  ** ******.*******:*****:*******

hLF      SRMDKVERLKQVLLHQQAKFGRNGSDCPDKFCLFQSETKNLLFNDNTECLARLHGKTTYE      660
bLF      SRSDRAAHVKQVLLHQQALFGKNGKNCPDKFCLFKSETKNLLFNDNTECLAKLGGRPTYE      658
         ** *:. ::********* **:**.:********:****************:* *: ***

hLF      KYLGPQYVAGITNLKKCSTSPLLEACEFLRK  691
bLF      EYLGTEYVTAIANLKKCSTSPLLEACAFLTR  689
         :*** :**:.*:************** ** :
```

**Fig. S7.** Sequence alignment of mature human (hLF) and bovine (bLF) lactoferrin. The alignment was prepared using Clustal O (1.2.4). * – identical, : – highly similar, . – similar
